# Habituation-Induced Expression of Immediate Early Genes in the Octopus Brain

**DOI:** 10.1101/2025.10.16.682951

**Authors:** Fabian Vergara-Ovalle

## Abstract

Understanding the neural basis of learning and memory in cephalopods remains challenging because of their behavioral complexity and the limited reproducibility of memory tasks. Here, I present a combined behavioral and molecular analysis using a simple habituation paradigm, Training Response to Artificial Prey (TRAP), in *Octopus bimaculoides*. Subjects showed a rapid, stimulus-specific decrease in exploration of a neutral prey-like object across trials. Habituation occurred in 100% of two- and six-month-old subjects, and multiple daily trials induced long-term memory lasting up to five days. Using whole-mount Hybridization Chain Reaction (HCR), I detected spatial activation of candidate immediate early genes, *creb* and *egr1*, during early memory consolidation. 15 minutes after learning, both genes were upregulated across the central brain, mainly in the subesophageal lobes, and axial ganglia. At 30 minutes, expression spread to the supraesophageal region, notably the vertical lobe, consistent with regions implicated in long-term memory. These results demonstrate a sequential activation from arms to subesophageal and supraesophageal regions, supporting proposed models of distributed information processing in the octopus nervous system.

**HIGHLIGHTS:**

- Training Response to Artificial Prey (TRAP) is a simple and highly repeatable method to evaluate memory in octopuses.
- Repeated trials induced robust, long-term habituation lasting up to five days.
- Immediate early genes *creb* and *egr1* were rapidly upregulated after training.
- (d) Gene expression spread sequentially across nervous system regions recruited for memory formation.

## INTRODUCTION

Cephalopods, particularly cuttlefish and octopuses, are invertebrates that exhibit a wide repertoire of behaviors. Many of these behaviors highlight their relatively advanced cognitive abilities when compared to other invertebrate groups (Mather & Alupay, 2016). Among these, memory has been the most extensively studied; different forms of memory in octopuses have been assessed, most often indirectly through observed learning processes. Octopuses have been shown to perform well in passive avoidance tasks (Shomrat et al., 2008), navigate mazes (Gutnick et al., 2020; Moriyama & Gunji, 1997), learn to solve problems with increased efficiency, such as opening jars or manipulating objects to the correct orientation (Richter et al., 2016; Fiorito et al., 1990), and form associations between visual cues and reinforcers (Bublitz et al., 2021; Bublitz et al., 2017; Sutherland, 1960; Boycott & Young, 1958). For a comprehensive review of octopus learning and memory, see Jozet-Alves et al. (2023). Despite the compelling body of evidence supporting memory in octopuses, developing efficient and repeatable paradigms for studying it remains a major challenge (Vergara-Ovalle et al., 2023; Boal, 1996). Octopuses frequently fail to respond to the experimental stimuli, due to low motivation and the lack of suitable reinforcers. To address this limitation, several behavioral paradigms that do not rely on reinforcement, and instead take advantage of the octopus’s instinctive responses to various stimuli, have been proposed, including novel object recognition (Vergara-Ovalle et al., 2023), spatial memory tasks (Hamlett, 2024), and episodic-like memory tasks (Poncet et al., 2022).

Despite the advances, these tasks are often difficult to replicate, and not all species, or even all subjects within the same species, are capable of performing them. This limitation results in experiments being constrained to the few animals that successfully complete the task, while excluding those belonging to species or groups unable to do so. Such constraints pose a significant challenge when aiming to investigate the neurobiological mechanisms underlying memory, since an ideal task should be easy to perform, rapid, repeatable, comparable between different species, and reliably executed by nearly 100% of the experimental subjects, as smaller sample sizes lead to less precise and accurate reliability estimates (Williams et al., 2025). In addition, valuable resources are expended when many subjects ultimately have to be excluded from the final analyses. From both a practical and ethical standpoint, behavioral tasks that can be performed by a greater proportion of subjects are therefore preferable, as they increase experimental efficiency and adhere to the principles of the 3Rs, which emphasize the refinement of methodologies to maximize the contribution of each animal used in research (Fiorito et al., 2014). Considering these requirements, I proposed the use of a simple, non-associative habituation learning paradigm as a tool to evaluate memory and its neural mechanisms in octopuses without the need for reinforcement.

Previous research has shown that lobes within the supraesophageal region are required for memory formation, including the Vertical lobe (Shomrat et al., 2008), Superior Frontal lobe (Boycott & Young, 1955), and the Inferior Frontal and Buccal lobes (Wells, 1965). However, most of these studies relied on generalized lesions of the region to assess their effects on memory, leaving the specific neural circuits and cell types involved unresolved. To investigate these processes in greater detail, one option is to employ molecular markers that increase in amount during neuronal activity following exposure to stimuli. Among such markers there are some genes which rapidly increase transcription after neuronal activation and are strongly associated with long-term memory consolidation, called Immediate Early Genes (IEGs) (Okuno, 2010), though IEGs have not been reported in octopuses prior to this study. Nevertheless, genes such as *creb* (cAMP response element binding gene), a stimulus induced-transcription factor (Lonze & Ginty, 2002), have been extensively studied in another mollusk widely used for memory research, *Aplysia californica* (Bonnick et al., 2012; Dash & Moore, 1996), and a homolog of *creb* has already been characterized in *Octopus vulgaris* (Sirakov et al., 2009). Similarly, *egr1* (Early growth response 1), another inducible transcription factor conserved as an IEG in vertebrates (Burmeister & Fernald, 2004), was recently reported to be abundantly expressed in the developing octopus brain (Styfhals et al., 2022).

However, the spatial expression of these genes during memory formation in cephalopods remains unknown. Identifying these patterns will reveal the neural structures and networks essential for memory formation in octopus, thereby enabling more precise comparisons of memory-related network properties across vertebrates and invertebrates.

In this study, I combined a novel behavioral habituation task in *Octopus bimaculoides* with whole-mount Hybridization Chain Reaction (HCR) to identify brain regions potentially involved in memory-related gene activation. This study contributes a new experimental model to explore the neural basis of memory in octopuses, offering a simple and reproducible task that can be combined with gene expression analyses to advance the study of animal cognition.

## METHODS

### Animals and Behavioral Task

For the behavioral task, *O. bimaculoides* subjects fell into three age groups: 2 months (n = 5), 6 months (n = 5), and 12 months (n = 2), which were tested in the habituation paradigm, hereafter referred to as TRAP (*Training Response to Artificial Prey*). The octopuses were obtained from the Cephalopod Breeding Center at the Marine Biological Laboratory (MBL), in Woods Hole, Massachusetts, USA. Animals were allowed a 24 h acclimation period and were housed in 80 L tanks with artificial seawater (35 ppt salinity) under suitable conditions, as described by Forsythe & Hanlon (1988).

All procedures were conducted in accordance with institutional guidelines for the care and welfare of laboratory animals and were approved by the Marine Biological Laboratory Institutional Animal Care and Use Committee (IACUC Protocol 24-07B).

As most benthic octopuses prefer crustaceans as prey (Markaida, 2023; Halil, 2019; Ambrose, 1984), I used a 3D-printed, odorless, resin crab: a stimulus that reliably elicited a prey-capture response (Fig S1). The fake prey size was matched to the corresponding group: 15mm, 45mm and 95mm for the two-, six- and 12-month animals, respectively. During each session, animals were exposed to six stimulus presentations (trials), spaced three minutes apart (Fig. 1a).

**Figure 1.**
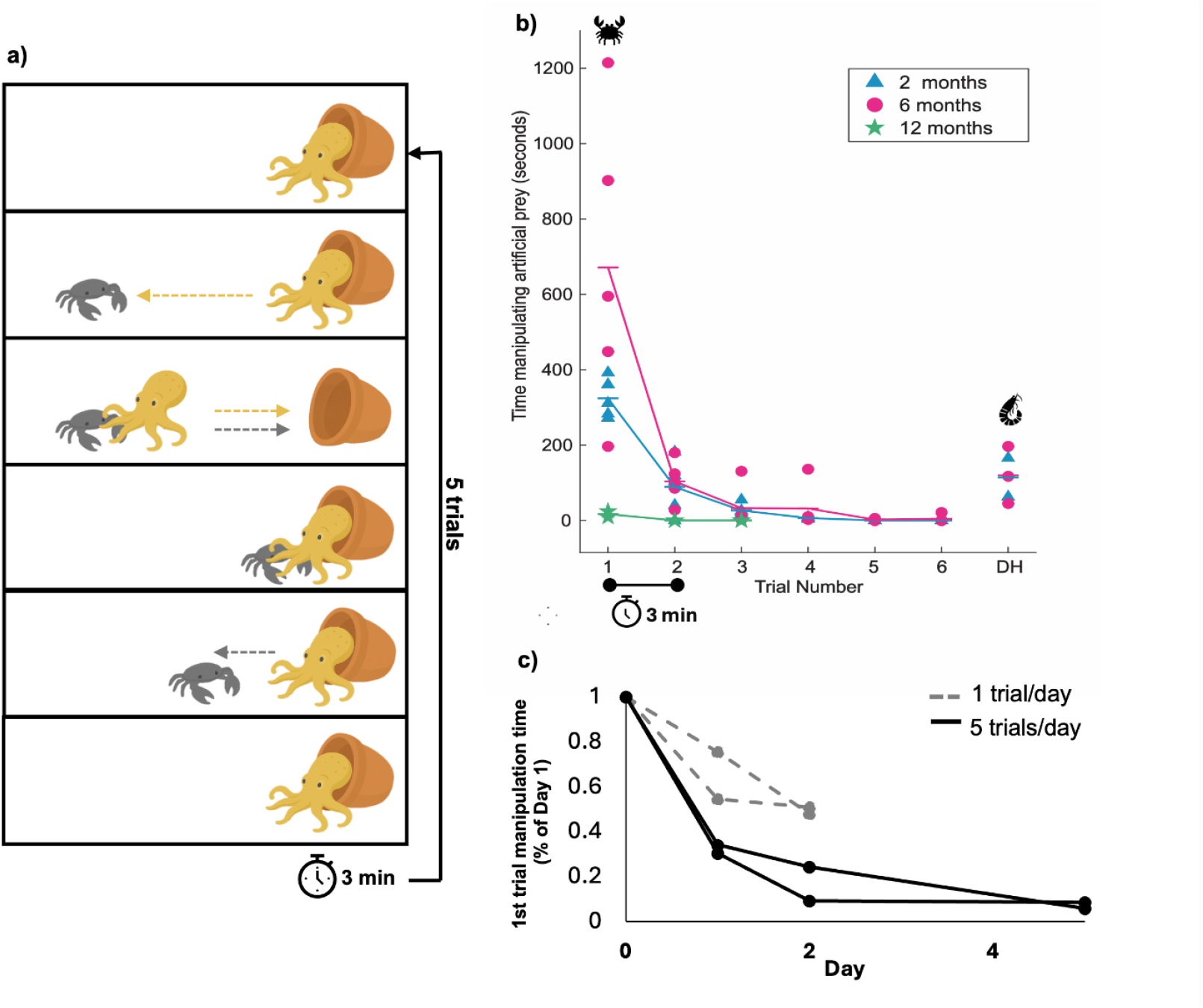
Octopuses rapidly habituate to artificial prey and form long-term memory with repeated exposure. (a) Schematic representation of the habituation task. An artificial prey-shaped plastic object was presented to the octopus, allowing manipulation, and the manipulation time was recorded. After the animal released the object, a 3-min interval was given before representation. (b) Graph showing habituation curves in 2- and 6-month-old subjects. Both groups exhibited a significant decrease in manipulation time after the first trial, with the response nearly absent by the fifth trial. Dishabituation occurred when a novel prey-shaped object was introduced. DH, dishabituation. (c) Memory retention tests. A single trial was sufficient to reduce the response after 24 h; however, repeated training (5 trials/day) induced long-term memory, with the reduced response persisting up to 5 days.

The total time the octopus spent physically manipulating the crab during each trial was recorded as a metric for engagement. The animals attacked the figure, grasping it with their arms and occasionally carrying it back to their dens, before eventually releasing and discarding the artificial prey. The fake crab was dropped from the water surface, landing at an approximate distance from the den of 7 cm for two-month-old subjects, and 35 cm for subjects six months and older. Notably, due to typical early-stage behavior, where young octopuses do not attack but rather consume prey that approaches or enters their den, it was sometimes necessary to place the fake prey adjacent to the den. Long-term memory was assessed by testing two groups of octopuses: one received a single trial per session, while the other received five trials per session. Each trial followed the same habituation protocol described before. Both groups were trained for one day, with one session per day. Memory retention was evaluated at 24 h, 48 h, and 120 h after the initial session, using the same behavioral metric of manipulation time. To assess dishabituation, the exploration time was measured for a novel object: a 3D-printed, resin shrimp. This object was similar to the original stimulus in size, texture, and color, differing only in shape.

### Tissue Collection and Gene Expression Analysis

To assess gene expression following behavioral habituation, five additional two-month-old subjects were analyzed using the TRAP protocol. Plus, a control group consisted of two subjects that freely explored objects without repetition and were euthanized 30 min after exploration. Subjects were anesthetized by immersion in 4% magnesium chloride solution in artificial seawater (8°C). Tissue samples were collected at 15 min (n = 3) or 30 min (n = 2) after the fifth trial, which marked the end of the acquisition phase. In this study, “acquisition” refers to the initial phase of the memory process, during which the behavioral response decreases progressively across trials but has not yet reached its minimum. During this period, the animal continues to acquire information from the environment through experience, a process interpreted as learning. Acquisition is considered the first stage of memory formation followed by consolidation, for the case of long-term memories. This conceptualization of acquisition has been widely discussed in the learning and memory literature (e.g., Noyes et al., 2021). In the present paradigm, acquisition was operationally defined as lasting until the response stabilized at its minimum value. Octopuses reached this minimum by the fourth trial, which was therefore taken as the end of acquisition; subsequent retention intervals of 15 or 30 min were measured from this point onward. Animals were fixed whole, with a small (~3 mm) incision in the mantle to facilitate fixative penetration in 4% paraformaldehyde (PFA) for 8 h, followed by 4% PFA + 30% sucrose for 24 h. Finally, tissues were transferred to 30% sucrose and either sectioned immediately or stored at –20 °C for later histological processing.

The protocol as described by Lovely et al. (2022) was followed with minor modifications. Briefly:

#### Dehydration

Tissues were sequentially immersed in increasing ethanol concentrations (50%, 70%, 90%, 100%; 5 min each). Excess ethanol was removed from the slide, and tissues were washed twice with 2× SSCT for 5 min.

#### Pre-hybridization

50 μL of hybridization buffer was added for 15 min.

#### Rehydration/Probe hybridization

Hybridization buffer was removed, and 50 μL of probe solution was added overnight at 37°C.

#### Wash

Probe solution was removed, and 50 μL of wash solution was applied three times for 15 min each at 37°C.

#### Amplification

Sections were washed with 5× SSCT for 15 min at 37°C and again for 5 min at room temperature. Fifty microliters of amplification buffer were added for 30 min at room temperature. Fifty microliters of hairpin solution were then applied for 6 h at room temperature in a humid chamber protected from light (omitted for negative controls).

#### Wash and Mounting

Hairpin solution was removed, and sections were washed twice with 50 μL of 5× SSCT for 30 min at room temperature. Samples were mounted using Vectashield antifade mounting medium with DAPI (Sigma Aldrich, MBD0020).

Using the published *O. bimaculoides* genome (Albertin et al., 2015), BLAST searches were performed to identify *creb* and *egr1* sequences. Based on these sequences, probe pools for HCR were designed for each gene (Table S1). Forty probe pairs were selected for *creb* and twenty for *egr1*, obtained as an oPool from Integrated DNA Technologies. oPools contained 50 pmol of each oligonucleotide in a dry pellet.

I targeted transcripts of *creb* and *egr1*, both candidates for activity-dependent transcriptional responses. Two types of controls were included: (1) animals allowed to explore freely without habituation, and (2) procedural controls processed without probe pools for *creb* and *egr1*. Signal detection and imaging were conducted using standard HCR v3.0 protocols. All solutions and buffers were obtained from Molecular Instruments. Images were acquired with a Leica Stellaris 8 STED microscope and analyzed using LAS X 4.7.0.28176 software and the open-access program Fiji (ImageJ).

### Statistical Analysis of TRAP

Behavioral responses during the TRAP paradigm were analyzed using the Wilcoxon signed-rank test to compare the duration of manipulation between trials.

## RESULTS

### Training Response to Artificial Prey (TRAP)

To establish a standardized task suitable for subsequent gene expression experiments, I evaluated the tactile response of octopuses to repeated presentations of a plastic crab to assess habituation (Fig. 1a). Both two-month-old and six-month-old subjects exhibited a statistically significant decrease in responsiveness to the neutral stimulus starting from the second trial (Wilcoxon signed-rank test, p < 0.05), with minimal exploration by the fourth trial. This reduction in response intensity was maintained throughout the remaining trials (Wilcoxon signed-rank test, p > 0.05). When a novel prey item (shaped like a shrimp) was presented at the end of the paradigm, subjects displayed a renewed increase in exploratory behavior, indicating dishabituation (Wilcoxon signed-rank test, p < 0.05). In contrast to the two younger groups, octopuses older than 12 months showed only minimal interest in the fake prey. In the first trial, they left their dens, approached the artificial crab, and lightly touched it, but did not use their suckers to manipulate it. In the following trial, their response diminished further, as they no longer emerged from their dens (Fig. 1b). Due to the weakness of these responses, subsequent analyses focused on the younger individuals, which displayed clearer behavior patterns.

One objective of this study was to identify IEGs in the octopus, which are overexpressed genes during the early stages of memory consolidation; that is, in the first minutes or hours after the acquisition phase. This consolidation phase leads to the formation of long-term memory. Therefore, in order to find IEGs, it is necessary that the memory being studied is maintained over the long term (above 24h). To this end, tests were conducted at 24, 48, and 120 hours post-acquisition, with either one or five trials per session. These tests revealed that even a single session is sufficient to induce a reduction in the long-term response (24h). However, this reduction was only about 30%, whereas a protocol of five trials per session resulted in a 65% decrease in response duration, which persisted up to 120 hours post-acquisition (Fig. 1c).

### Early consolidation involves distinct brain and arm activation of *egr1* and *creb*

To identify brain regions involved in the formation of habituation memory in octopuses, I performed HCR to assess the expression of *creb* and *egr1*. Samples were taken 15 and 30 minutes after the fourth trial on the first day of the habituation task, corresponding to the beginning of the memory consolidation phase. 15 minutes after the acquisition phase, both *egr1* and *creb* showed increased expression across the central brain compared to controls. The highest levels were observed in the subesophageal region, including the palliovisceral, pedal, and brachial lobes (*egr1*: 16.98% vs. 0.22%; *creb*: 11.51% vs. 3.65%), as well as in the anterior basal lobe of the supraesophageal region (*egr1*: 3.07% vs. 0.08%; *creb*: 1.54% vs. 0.73%) (Fig. 2b,e).

**Figure 2.**
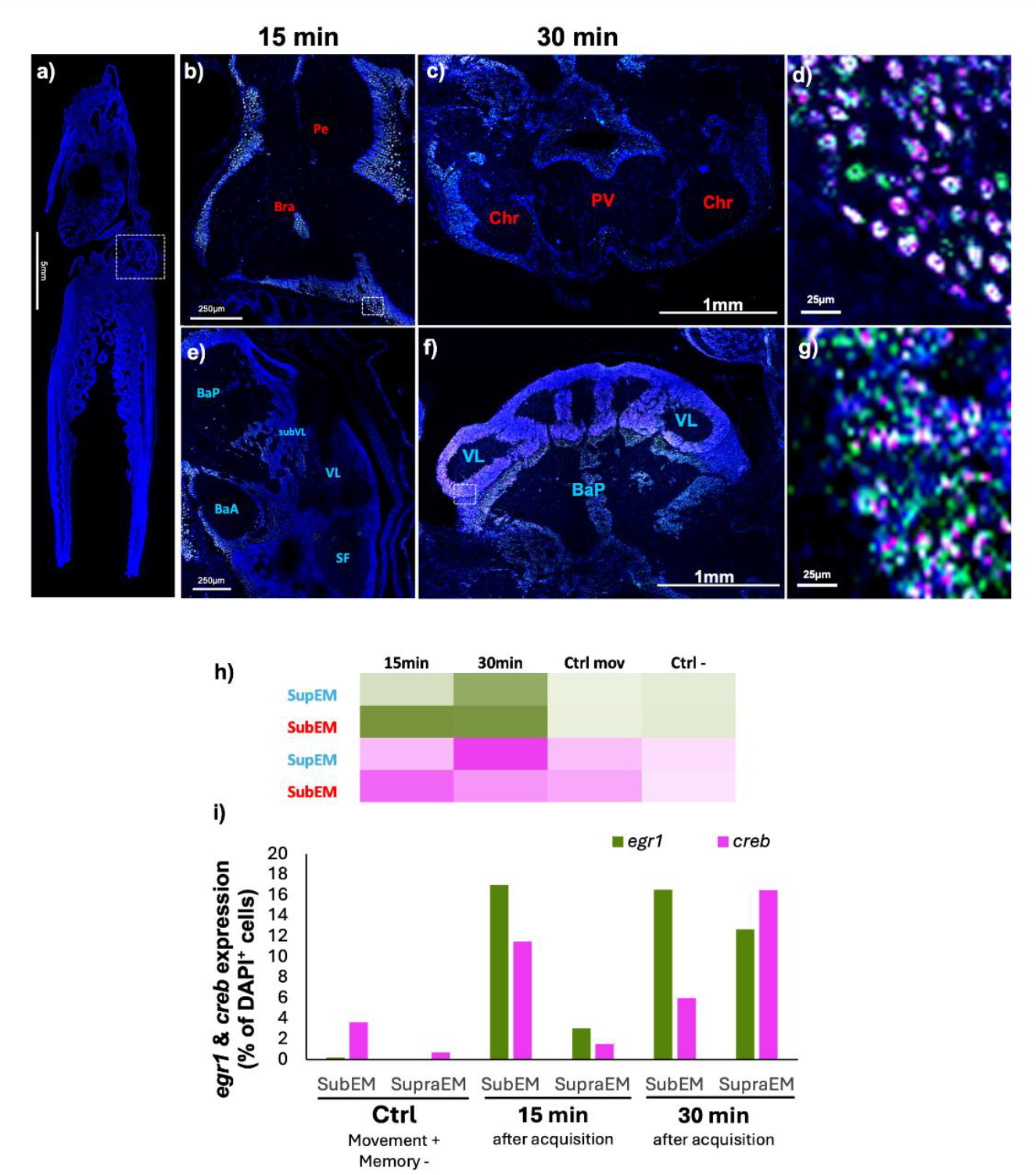
Spatiotemporal recruitment of *creb* and *egr1* in the octopus brain during early memory formation. (a) Sagittal section of *O. bimaculoides* stained with DAPI, highlighting the central brain (white box). At 15 min, strong expression of *egr1* (green) and *creb* (pink) is observed in the subesophageal mass (SubEM) (b), while the supraesophageal mass (SupraEM) shows low activity (e). Panel (d) shows a magnified view of the boxed area in (b). At 30 min, *egr1* expression remains elevated in the SubEM whereas *creb* decreases (c); in contrast, both genes show a marked increase in the SupraEM (f). Panel (g) shows a magnified view of the boxed area in (f), with granular cells. (h) Heatmap showing *egr1* and *creb* expression in SubEM and SupraEM across groups. (i) Percentage of DAPI-positive cells expressing *egr1* and *creb*. Pink regions indicate *creb* expression, and green regions indicate *egr1* expression. Images acquired with 20x.

At 30 minutes, *egr1* expression remained elevated in the subesophageal region (16.5%) and increased in the supraesophageal region (12.7%), particularly in the vertical and posterior basal lobes. In contrast, *creb* expression decreased in the subesophageal region (6%) but increased in the supraesophageal region (16.5%), mainly in the vertical lobe (Fig. 2c,f).

In addition, both genes showed increased expression in the arms 15 minutes after the habituation task (*egr1*: 2% vs. 14%; *creb*: 15% vs. 38%). This expression was detected in the axial ganglia and along the entire length of the arm (Fig. 3).

**Figure 3.**
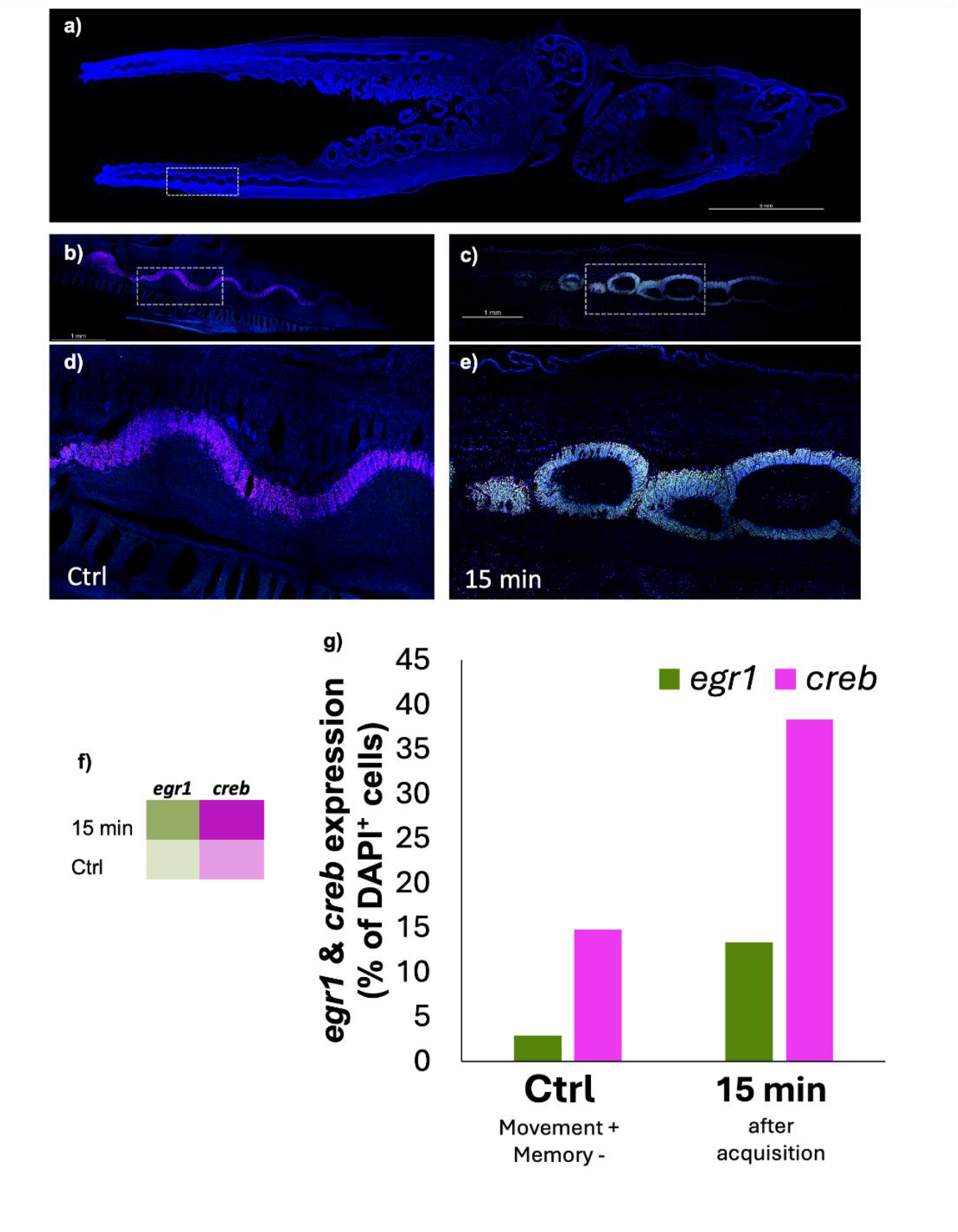
Peripheral neural circuits in the arms participate in sensory processing and memory after habituation. (a) Sagittal section of whole *O. bimaculoides* stained with DAPI, showing the nervous system and highlighting part of the axial ganglia within the arm (white box). Expression of *egr1* and *creb* is observed in the axial ganglia of the arms in the control group (b, d) and 15 minutes after TRAP (d, e). (f) Heatmap representing the expression of each gene across groups. (g) Percentage of DAPI-positive cells showing expression of *egr1* and *creb*. Images acquired with 20x.

## Discussion

In this study, I present a non-associative, habituation learning task for octopuses that can be used to trace the neural mechanisms of learning in the octopus nervous system. The paradigm relies on repeated presentations of an artificial crab, one of the octopus’ preferred prey items. Learning was demonstrated by initial, tactile interest in the fake prey item, followed by reduced interaction time in subsequent trials. Once the octopuses learned that the prey was inedible and unusable during the first trial, engagement significantly dropped. The octopuses were dishabituated with the introduction of a novel fake prey item, confirming a learning of stimulus specificity.

Crabs are among the most common and preferred prey items for many octopus species, which makes them an ecologically meaningful stimulus in behavioral experiments. Their strong ecological relevance explains why artificial crabs have already been used in different contexts, such as assessing welfare through enrichment and health status evaluations (Borrelli et al., 2020; Amodio, 2014) or even as decoys in fisheries (Markaida et al., 2019). Presenting such prey-like stimuli in controlled conditions increases the likelihood that the observed responses recruit the same mechanisms involved in natural predator–prey interactions. Moreover, tasks grounded in ecologically relevant cues may reduce the stress associated with exposure to arbitrary or unfamiliar conditions, thereby generating behavioral data that more faithfully reflect the animal’s natural behavior repertoire. Within this framework, this study introduces a novel habituation task in octopuses using artificial prey, extending an approach that has been widely applied across other animal models. The ecological relevance of the task ensured consistent engagement, providing the basis for the high success rate observed: 100% of two- and six-month-old subjects demonstrated full learning from the second trial onward. This is a level of task completion rarely achieved in octopus behavioral studies, where a substantial proportion of animals often fail to complete the task and are excluded from analysis (Poncet et al., 2022; Bublitz et al., 2021; Richter et al., 2016).

Expanding on the time scale of the experiment, octopuses were able to form long-term memories 24 hours after training with exposure to either a single trial or five trials per day. However, the animals were far less willing to explore the object after they had been exposed to five trials rather than just the single trial. This relationship between trial number and memory strength aligns with findings in other molluscan models. In *A. californica*, long-term sensitization is approximately five times stronger when animals receive four trials per day compared to a single one (Frost et al., 1985). This effect likely occurs because a single event is sufficient to initiate memory consolidation; that is, the process by which transient experiences are stabilized into more durable memory traces (Squire et al., 2015). In octopuses, the ability to consolidate information from a single encounter may be especially relevant given the ecological importance of rapid learning in predator– prey interactions (Hanlon, 2024; Hanlon & Messenger, 2018). At the same time, repeated exposures are often required to strengthen these traces through more enduring forms of synaptic plasticity, providing a neural basis for memories that are both more robust and persistent (Goto, 2022).

While the younger groups of octopuses behaved very similarly, octopuses older than one year would visually inspect the prey item before returning to their den, without making meaningful tactile contact. The observed reduction in fake prey manipulation may reflect a form of behavioral inhibition, defined here as the suppression or withholding of tactile exploratory behavior. Such inhibition suggests that adult octopuses visually identified the stimulus as neutral and therefore reduced unnecessary tactile exploration. This age-related improvement in visual recognition is similar to what has been previously observed in adult *O. maya* during the Novel Object Recognition task (Vergara-Ovalle et al., 2023), where four-week-old octopuses were unable to discriminate familiar from novel objects, unlike older subjects. This developmental trajectory is likely associated with the progressive maturation of the octopus nervous system throughout its lifespan (Vergara-Ovalle et al., 2022).

This study provides evidence linking the behavioral habituation paradigm directly to spatial activation of candidate IEGs in the octopus nervous system. IEGs such as *creb* and *egr1* are rapidly and transiently expressed in neurons following synaptic activity and have long been used as molecular markers of learning and memory processes. In this case, increased in-situ expression of *creb* and *egr1* appeared in the early phases of long-term memory consolidation, specifically in the brain and the axial ganglia. At 15 minutes post-acquisition, increased expression of both genes was detected throughout the central brain, predominantly in the subesophageal region: the palliovisceral, brachial, and pedal lobes. The subesophageal region has been compared to the vertebrate cerebellar stem and spinal cord, as it is involved in controlling arm and body movements necessary for exploration. The region has both afferent and efferent fibers from the arms, mantle, eyes, mouth, and olfactory organ, as well as a high number of interneurons that filter and regulate information (Shigeno et al., 2018). These results suggest that during the first minutes after the acquisition phase, the subesophageal region, an area involved in controlling arm and body movements necessary for exploration, is selectively activated, as indicated by the higher expression of *egr1* and *creb* compared to control animals that explored the object freely without repeated exposures. This early activation likely reflects the engagement of motor and sensory circuits that are critical for encoding the habituation memory.

At 30 minutes post-acquisition, increased *egr1* and *creb* activity was additionally observed in the supraesophageal region, with strong labeling in the vertical lobe. This region is critical for long-term memory formation in octopuses, as demonstrated by previous research: after ablation of the vertical lobe, octopuses could learn tasks, but memory did not persist beyond 24 hours (Young, 1960). Similarly, Shomrat et al. (2008) observed that octopuses required more trials to suppress attack behavior when the vertical lobe was tetanized. Given its function and connectivity, the vertical lobe has been compared to the mushroom bodies in arthropods and the hippocampus in vertebrates (Shomrat et al., 2015). The overexpression of *egr1* and *creb* in a time dependent manner suggests that the vertical lobe, already known from classical experiments to be essential for long-term memory, is indeed a site of rapid gene activation during memory consolidation, linking molecular activity to established behavioral outcomes.

Additionally, increased expression of both genes was observed in the axial ganglia neurons of the octopus arms 15 minutes after the task. *egr1* had similar expression levels to the subesophageal region, whereas *creb* levels were more than double what was expressed in the subesophageal region at the same time point. These inferred patterns of neuronal activity align with previously described sensorimotor models for the octopus, in which sensory information flows from the arms to the central brain, first through the subesophageal region and then to the supraesophageal region, with staged processing occurring at each level, rather than a single centralized processing center as in vertebrates (Hochner et al., 2023; Hooper, 2020). This pattern of gene expression suggests that the arms are not merely peripheral effectors but actively contribute to early stages of sensory processing and memory-related activity. Such local processing provides a distributed substrate that supports exploratory and learning behaviors in octopuses. Importantly, this arm-based activity likely interacts with central brain circuits, allowing integration of the information from different senses, and stabilized into long-term memory representations. More broadly, studies like this one reveal the structure and timing of engaged neural circuits, enabling comparative analyses of brain function in different nervous systems that may contribute to our understanding of the evolutionary origins of complex behaviors (Binyamin Hochner et al., 2006).

The present study establishes a reliable habituation task for octopuses that is simple to implement and effectively induces short and long-term memory. It provides evidence that *creb* and *egr1* are rapidly and selectively activated across distributed regions of the octopus nervous system, including both the brain (the subesophageal and the supraesophageal regions) and the arms, during early memory consolidation. These results reveal a time and spatially organized network of neural structures underlying memory formation, demonstrating that both central and peripheral circuits contribute in a time dependent manner. Together, the TRAP paradigm and the mapping of immediate early gene activation provide a powerful framework for dissecting the neural basis of cognition in cephalopods, that can be expanded upon to understand the flow of information and memory consolidation in freely behaving animals.

## ACKNOWLEDGEMENTS

The author gratefully acknowledges the Grass Foundation for its support in carrying out this project, and Kristianna Lea for her helpful commentary on the manuscript. The author also thanks all members of the Grass Lab for their guidance, technical assistance, and stimulating discussions throughout the project, as well as the administrative staff at the Marine Biological Laboratory for their invaluable logistical support.

## GRAPHICAL ABSTRACT

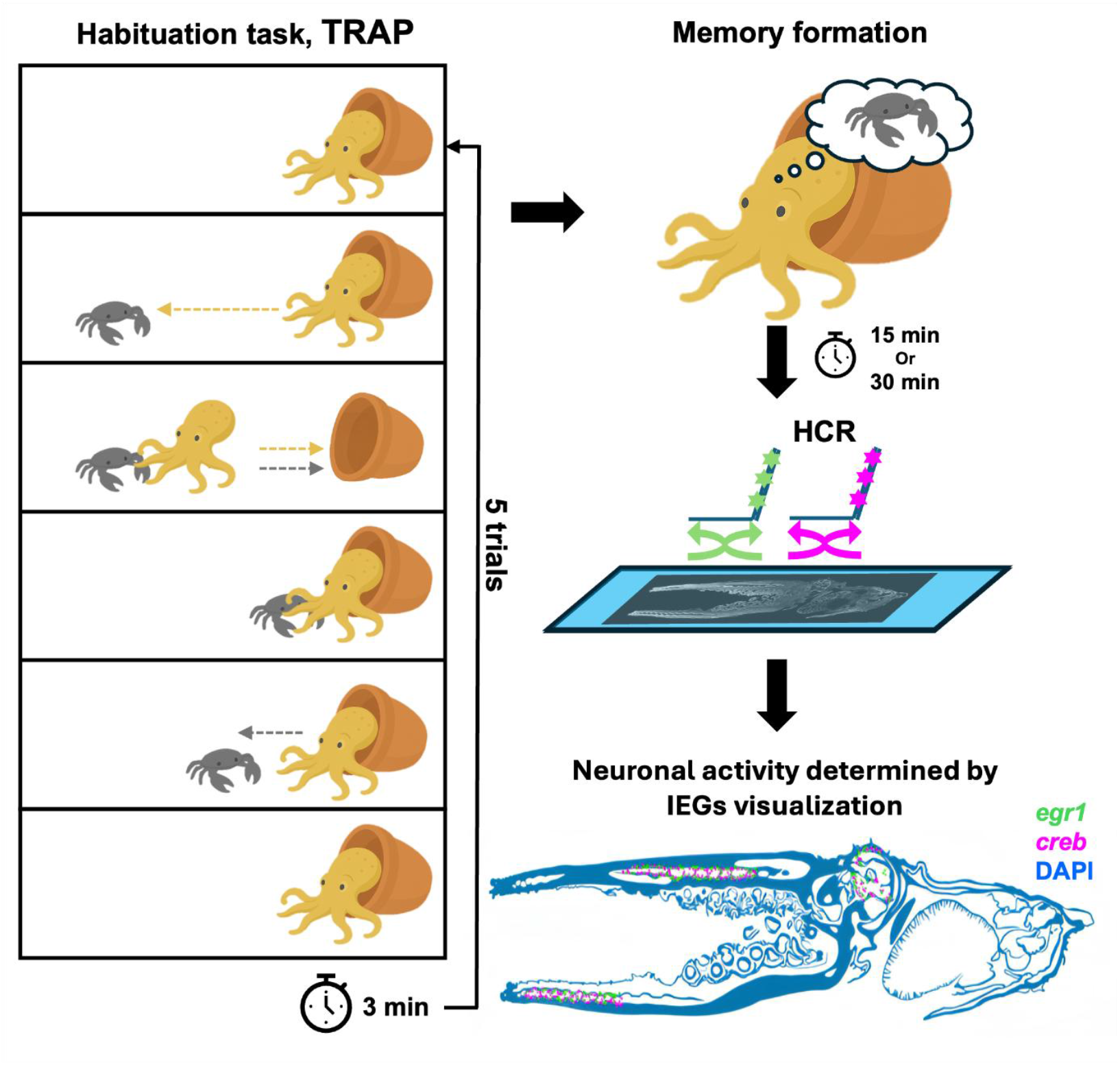

