## Supplementary Figure 1 for "Habituation-Induced Expression of Immediate Early Genes in the Octopus Brain"

**Supplementary Figure S1**

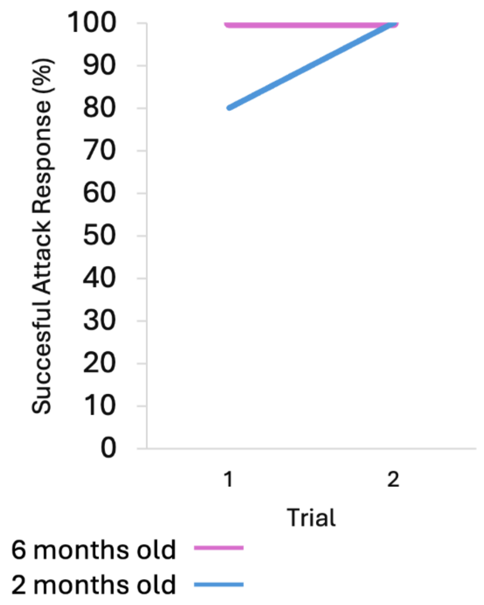


**Supplementary Figure 1**. Percentage of successful attack responses across two consecutive trials in octopuses tested with the Training Response to Artificial Prey (TRAP) paradigm. Animals were presented with a false prey stimulus at 2 months of age (magenta, n = 5) and 6 months of age (teal, n = 5). Trials were separated by a 3-minute interval.
